# Dissociating instructive from permissive roles of brain circuits with reversible neural activity manipulations

**DOI:** 10.1101/2023.05.11.540397

**Authors:** Daniel Quintana, Hayley A. Bounds, Jennifer Brown, May Wang, J. Simon Wiegert, Hillel Adesnik

## Abstract

The brain has a remarkable capacity to recover from damage. Despite the value of such recovery, it has made determining which areas are critically involved in a specific brain function challenging. Different types of manipulations to brain tissue – from mechanical or chemical lesions to genetic ablations or pharmacological agents – often yield conflicting conclusions. In some cases, ablating a brain area causes only a temporary disruption because a downstream region can homeostatically recover from its loss of input, while in others it leads to permanent impairment due not to the damaged area per se, but because of the degeneration of distant areas. We sought to address these issues by exploiting the flexibility and reversibility of optogenetic manipulations. We applied a three-stage optogenetic approach to the mouse’s primary visual cortex (V1) in a contrast detection task, beginning with intermittent silencing, followed by sustained illumination across multiple days of task-related behavior, and concluding with a return to intermittent silencing. Initially, intermittent V1 inactivation strongly impaired task performance, while during persistent multi-session inactivation, the animals’ visual detection entirely recovered to pre-silencing levels. However, this recovery was itself reversible, as returning the animals to intermittent V1 inactivation reinstated the behavioral deficit. Suppressing activity in the superior colliculus (SC) concurrent with persistent V1 inactivation reversed the animals’ recovery, demonstrating that SC is a key area that drives behavioral compensation in the absence of V1 function, even while under normal conditions V1 activity is crucial. These results support a scenario where primary sensory cortex is in the default circuit mice use to detect visual stimuli, but in its absence, midbrain regions can compensate for it. More generally, this temporally controllable optogenetic perturbation paradigm could be useful to adjudicate whether a brain area instructs or is accessory to a specific brain function.

## INTRODUCTION

Neuroscientists rely on targeted perturbations to causally map functions in the brain^1^. Yet since the brain is highly interconnected, manipulation of one area can impact behavior through indirect effects on other brain regions, complicating the interpretation of such results^2,3^. Permanent lesions of one area can lead to dysfunction or degeneration of other brain regions, making it difficult to determine whether the functional consequences of the lesion are due to loss of the targeted region or to those that are indirectly damaged. Moreover, the often-observed recovery of behavioral performance after a permanent lesion can further complicate the interpretation of lesion experiments: either the lesioned area was never directly involved, or another brain area was able to compensate for its loss^4,5^. Compared to irreversible lesions, acute optogenetic manipulations avoid potential indirect neurodegenerative effects^2,6^, permit interleaved control trials, and can address the timescales for when neural activity in specific regions is necessary for behavioral output^7,8^. Yet, recent studies have highlighted how the results of acute and irreversible inactivation can conflict^4,5,9^. For example, acute inactivation of motor, hippocampal, or sensory areas in animals can lead to profound behavioral deficits, but animals with permanent lesions are either unaffected or can recover performance within just a few days^4,5,9,10^. In one striking example, acutely silencing a forebrain nucleus of the songbird severely degraded song production, but the deficit could be attributed to an indirect effect on activity in a downstream area^4^. After lesion, the downstream area homeostatically adjusted its activity and the animal’s behavior recovered. This led the authors to propose that a brain region can either be “instructive” – providing essential information that is used in the computation for a behavioral task, or “permissive” – acutely required to set the gain or sensitivity of another region, but not providing essential information or computation (Fig. 1a). Similarly, a study of a whisker-dependent task observed that acute silencing of barrel cortex impaired behavior, but animals with permanent barrel cortex lesions could completely recover performance after a single training session. This study offered two possible conclusions: either barrel cortex is instructive but is redundant with another area that compensates for its loss^5^ (Fig. 1b); or barrel cortex is permissive and acute inactivation transiently disrupts the instructive area. The inability to properly interpret the results of acute manipulations is a major roadblock in neuroscience.

**Figure 1.**
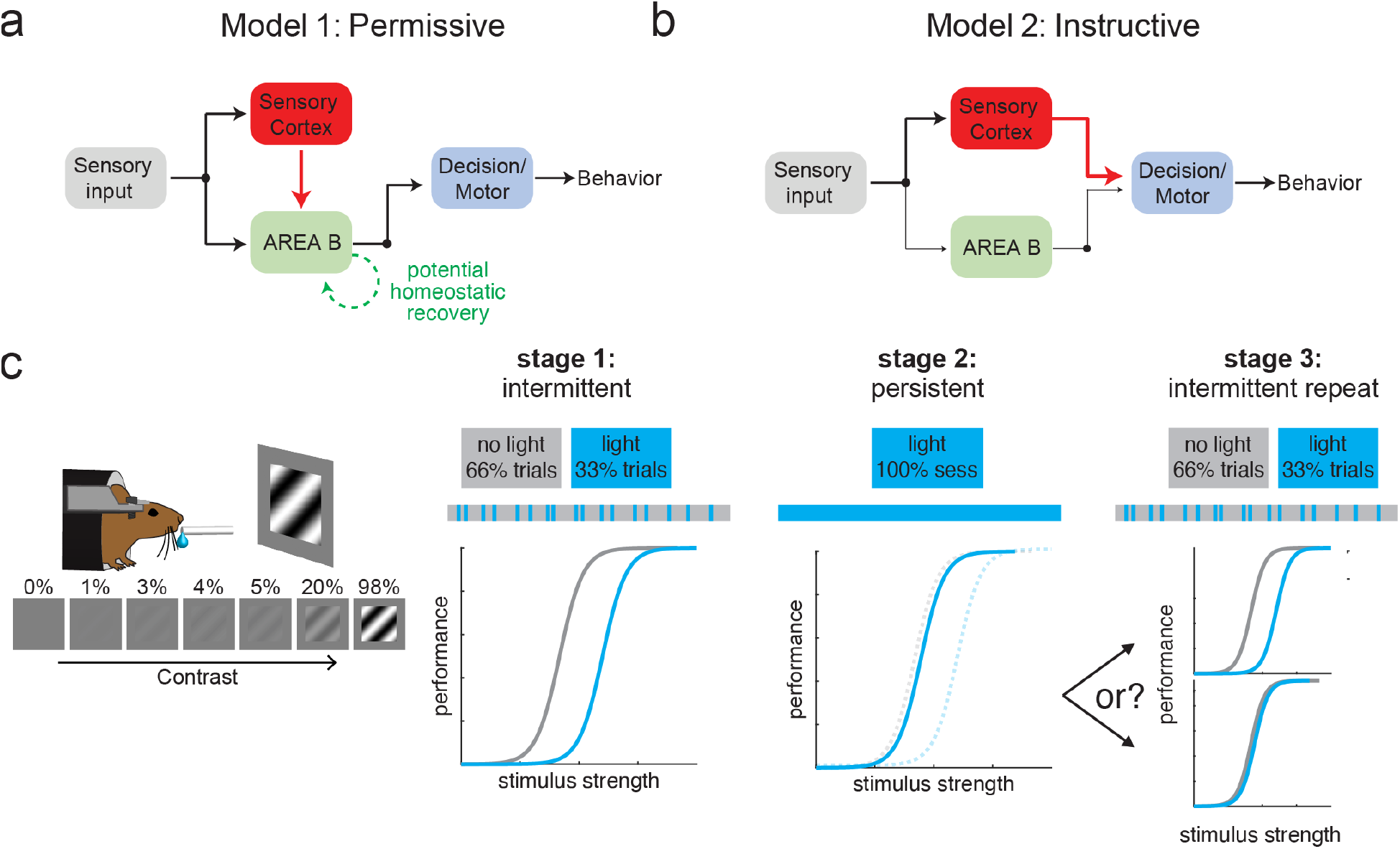
A three-stage optogenetic inactivation paradigm for dissociating instructive from permissive roles of a targeted brain circuit. **A)** A permissive model for the circuit. In this model, Area A is only needed to keep Area B in a functional state so that Area B instructs correct performance. When Area A is removed or permanently silenced (bottom), Area B can homeostatically recover (curved green arrow) from its loss of drive from Area A and restore correct behavior. **B)** An instructive model for the circuit. In this model, Area A directly instructs correct performance, but when it is removed or permanently silenced, a redundant accessory circuit through Area B restores performance (bottom). **C)** An approach to distinguish among these models with chronic but reversible inactivation: three successive stages of intermittent, persistent, and repeat intermittent silencing of the targeted circuit. Bottom: schematic plots of behavioral performance on a relevant task showing performance in each stage for light off (gray) and light on (blue) trials. For stage 2, the dotted lines are the schematized performance data from stage 1 replotted for comparison. In this example set of results, in stage 1, inactivation reduces performance, but in stage 2 performance recovers. In stage 3, intermittent inactivation either returns to reducing performance (top) or no longer has any effect (bottom).

Comparing lesion and acute silencing of visual cortex yields a similarly ambiguous picture of its role in visual perception. In humans, V1 lesions lead to permanent loss of visual perception, often in circumscribed regions of the visual field, while some residual visual function is often left intact^11^. In line with this, classic lesion studies in rodent V1 have shown that V1 appears dispensable for a range of visual function. After V1 lesion, rats rapidly relearned visual tasks and recovered much of their acuity^12^, and in mice V1 lesions spare detection of low spatial frequency or high contrast gratings^13,14^ but impair the detection of high spatial frequency gratings and may impair orientation discrimination (though studies conflict)^8,14^. However, a key complication for interpreting these results is that V1 lesions lead to degeneration of neurons in the primary visual thalamus, the lateral geniculate nucleus (LGN), which also projects to non-V1 areas. In one study, the severity of deficit after lesion depended on whether the lesioning method drove LGN degeneration, suggesting that LGN degeneration per se, and not the loss of V1, is what drove the visual deficits^15^. Conversely, acute optogenetic silencing, which does not cause distal degeneration, has been shown to affect, but not abolish, a range of visual abilities, including the detection and discrimination of orientation and contrast^7,8,13,16–20^, even at low spatial frequencies^7,13,16^.

Thus, the role of V1 in basic visual perception remains unresolved, and either V1 is not directly involved in basic visual perception, and/or it can be compensated for by other regions, such as the superior colliculus (SC), which is a major target of retinal and non-retinal visual inputs and has been implicated in visual tasks including contrast detection^21^. Because this prior work has not yielded a concrete picture of V1’s role in visual perception, we sought to address this ambiguity with an optogenetic approach that can be both reversible and sustained over different time scales.

We thus validated and employed a three-stage optogenetic approach to determine the role of V1 in the detection of visual stimuli (Fig. 1c). We found that intermittent V1 silencing strongly reduced a mouse’s ability to detect low contrast stimuli, consistent with prior reports. We then silenced V1 over the entire session for several days and found that mice almost completely recovered behavioral performance. Finally, we returned these fully recovered mice to intermittent silencing and found that the mice reverted to failing on optogenetic silencing trials. These data suggest that V1 is instructive in the visual perceptual task but can be compensated for by other brain areas in the continued absence of its normal activity levels. To identify a potential compensatory brain region, we optogenetically silenced the SC and found that this drastically reduced task performance in mice that had recovered from persistent V1 silencing. Taken together, these results demonstrate the remarkable flexibility of the brain to re-route activity to support behavior, demarcate the timescales and reversibility of these dynamic adjustments, and identify midbrain areas as key loci that can compensate for the loss of cortical function. This three-stage optogenetic approach of intermittent-persistent-intermittent silencing might be more broadly applied to better delineate whether a given brain area instructs or permits a specific brain function.

## RESULTS

### A flexible optogenetic strategy for both acute and persistent silencing of neural circuits

We reasoned that we could help address the ambiguities of existing perturbation approaches by employing a flexible optogenetic inactivation strategy in which one could acutely or chronically silence a brain area in the same animal. With such an approach, one could first measure the acute effects of inactivation of a brain area on a behavior, then chronically silence it to mimic an irreversible lesion, and finally return the brain area to acute inactivation (Fig. 1c). In cases where acute inactivation impairs behavior, animals might recover behavioral function during chronic optogenetic silencing, as they often do after irreversible lesions. In such cases, one might be able to address the apparent conflict by returning to acute, intermittent silencing in the recovered animals. This third test could be highly informative: if recovered animals no longer showed any effect of acute silencing, this would suggest that the brain region only played a permissive role in the initial condition (Fig. 1c). Conversely, if a return to acute silencing in recovered animals reinstates the behavioral deficit, the brain region is more likely to directly contribute to the computational process in an instructive manner. Finally, concurrent optogenetic suppression of other brain regions during the recovery phase could reveal the degree and locus of compensatory function.

To develop and leverage the proposed optogenetic strategy, we focused on V1, which is among the best studied areas in the mammalian brain, but whose role in different aspects of visual perception remains in doubt. To touch base with prior work, we first investigated the impact of permanent lesion on performance in a contrast detection task^13,14^. We trained mice to detect gratings of a spatial frequency that prior lesion work implies does not require V1, but prior work with acute manipulations implies should require V1, enabling us to investigate a key unresolved ambiguity^7,13,14,16^. We quantified performance using the detection threshold, or c50 (Methods). Lesion of V1 by aspiration caused an acute reduction in task performance, followed by a substantial, albeit incomplete, recovery over the subsequent week (Fig. 2a-b; Pre vs Day 1 (∼24 hours after surgery): p<0.05, Pre vs. Days 7-10: p<0.05, Day 1 vs Days 7-10: p<0.05; n=7 mice). There was a qualitative but not significant relationship between lesion size and initial deficit, and no correlation with the behavioral deficit after performance stabilized (Fig. S1e-f). To assess the extent to which surgery itself may have contributed to the observed deficits, we lesioned primary somatosensory cortex (S1) in a separate cohort of mice. Lesioning S1 had a much smaller effect on visual behavior (Fig. S1a-b), demonstrating that by 24 hours after lesion, the behavioral effect we observed with V1 lesions was due specifically to the loss of V1 and not due to surgery *per se* (Day 1 post lesion S1 vs V1: p<0.05, n=10 mice).

**Figure 2.**
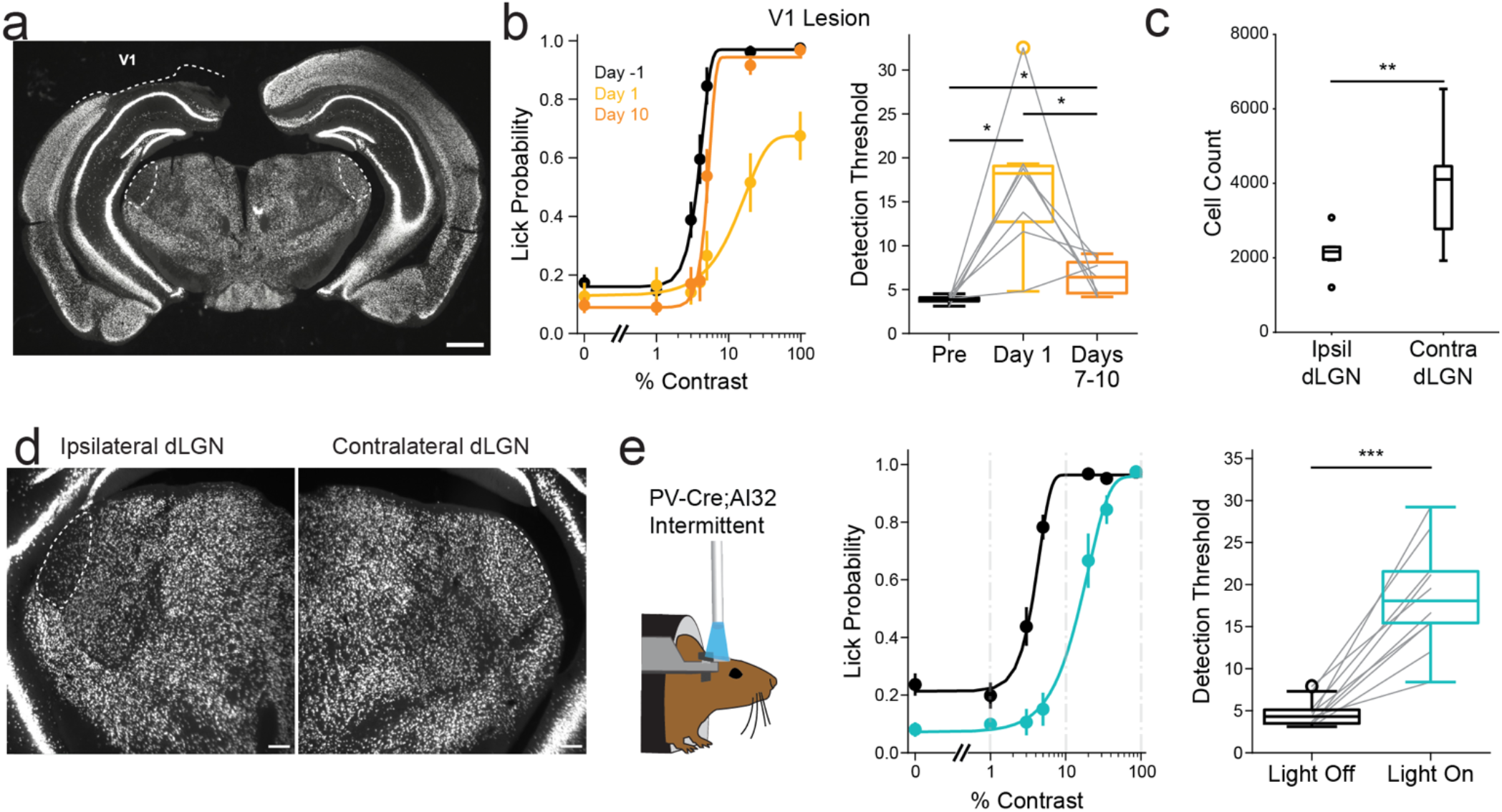
Conflicting effects of lesion and acute inactivation of V1 on behavioral performance in a visual detection task. **A)** Coronal section from a representative mouse stained with anti-NeuN, showing the extent of the V1 lesion. Dashed circles indicate the locations of the bilateral dorsal lateral geniculate nucleus (dLGNs). Scale bar: 1 mm. **B)** Left, psychometric performance curves for mice with V1 lesions measured on the day before lesioning (Day −1, black), the day after lesion (Day 1, yellow), and Day 10 post-lesion (orange). Lines represent Weibull function fits to the behavioral data. Right, boxplot showing detection thresholds averaged across six pre-lesion days, Day 1 post-lesion, and Days 7–10 post-lesion. Statistical comparisons: repeated-measures ANOVA, p < 0.05; paired t-tests with multiple comparison correction: Pre vs. Day 1: p < 0.05, Pre vs. Days 7–10: p < 0.05; Day 1 vs. Days 7–10: p < 0.05; n = 7 mice. **C)** Quantification of dLGN cell counts comparing ipsilateral (to the V1 lesion) and contralateral (to the V1 lesion) hemispheres (p < 0.01, paired t-test; n = 7 mice). **D)** Expanded view of the images of left (ipsilateral to the V1 lesion) and right (contralateral to the V1 lesion) dLGN from the section in (A). Scale bar: 100 μm. **E)** Left, psychometric performance curve for PV-Cre;Rosa-LSL-ChR2(Ai32) mice during intermittent silencing. Lines are Weibull function fits. Blue, light on; black, light off. Right, boxplot of detection thresholds during intermittent photostimulation; p<0.0001, paired t-test. n=10 mice.

Consistent with prior reports^12,13^, these data show mice exhibit substantial recovery of contrast sensitivity following V1 lesion. This raises two important questions: why does visual behavior recover at all and why is it incomplete? Behavior might recover because V1 is merely permissive and the brain homeostatically scales up the gain of the instructive area to compensate for V1’s loss. Alternatively, V1 is instructive, but behavior recovered because the animal learned to perform the task using alternative brain structures^12,13^. Recovery might be incomplete because either V1 is essential for a small component of behavioral performance, or lesioning V1 leads to degeneration of other visual areas that are. Indeed, inspection of the primary visual thalamus (LGN) in V1-lesioned mice showed a profound reduction in the cell counts ipsilateral to the lesion, consistent with a recent report (Fig. 2c-d; p<0.01, n=7 mice)^15^. Thus, based on these lesion results alone, we cannot conclude whether V1 is instructive or permissive in the contrast detection task, and we cannot rule out that distal degeneration of other visual areas is the source of the lasting behavioral deficit. These data highlight key problems with interpreting the results of conventional lesioning studies.

To begin to address the challenges of interpreting both the recovery after brain damage and any lasting effects, we sought to use a reversible perturbation to clarify the role of V1 in basic visual functions in mice, as other studies have done^13,16,17^. We silenced excitatory neurons by optogenetically stimulating GABAergic neurons, which caused a profound reduction in contrast sensitivity on illumination trials (Fig. 2e; PV-Cre; Rosa-LSL-ChR2(Ai32): p<0.0001, n=10 mice). Similar illumination in mice not expressing any opsin did not affect performance (Fig. S1c, p = 0.86, paired t-test, n=4 mice), controlling for any direct effects of the illumination on behavior. At face value, this acute inactivation experiment supports an instructive role of V1 in contrast detection, but it seemingly conflicts with the substantial recovery seen in mice with V1 entirely removed (Fig. 2b). Taken together, the results of irreversible and acute, reversible lesions of V1 highlight the difficulty of drawing definitive conclusions about its role in behavior even with a combination of approaches.

While optogenetic manipulations have conventionally been used to silence brain regions or cell types only briefly, we reasoned that prolonged optogenetic silencing might add crucial information by mimicking a permanent lesion, but ultimately still being reversible. For this approach we employed direct optogenetic silencing of cortical excitatory neurons by expressing the blue light silencer soma-targeted GtACR2 via the construct somBiPOLES^22^, which provides membrane-targeted expression that can be difficult to achieve when expressing GtACR2 alone^23^. To test the efficacy of silencing neural activity with somBiPOLES under both intermittent and persistent illumination conditions, in non-behaving mice, we recorded neural activity with multielectrode arrays across the layers of V1 while illuminating the cortex briefly on every third trial, and then persistently for 40 continuous minutes (2mW/mm^2^, 455 nm) (Fig. 3a-c). In somBiPOLES-expressing mice, brief illumination resulted in immediate and strong suppression of both spontaneous and visually evoked neural activity (Fig. 3c-e; 83±4% reduction at 100% contrast, p<0.05, Wilcoxon signed rank test). Next, we asked if uninterrupted sustained illumination could effectively silence cortical activity for the entire duration of the illumination, and whether neural activity would rapidly recover after cessation of illumination to rule out potential photo-toxic effects. Indeed, during persistent illumination, the mean firing rate averaged over all recorded units was suppressed by 76±4% (Fig. 3d, e; at 100% contrast, p<0.05, Wilcoxon signed rank test). After the cessation of sustained illumination, the mean activity was indistinguishable from baseline levels (Fig. 3e; p=0.53, Wilcoxon signed rank test), arguing against potential issues with prolonged illumination^24^ at this light level, 2mW/mm^2^, or that prolonged flux of ionic currents might lead to cellular alterations^25^. These experiments confirm the efficacy, stability, and reversibility of these acute and more persistent optogenetic manipulations.

**Figure 3.**
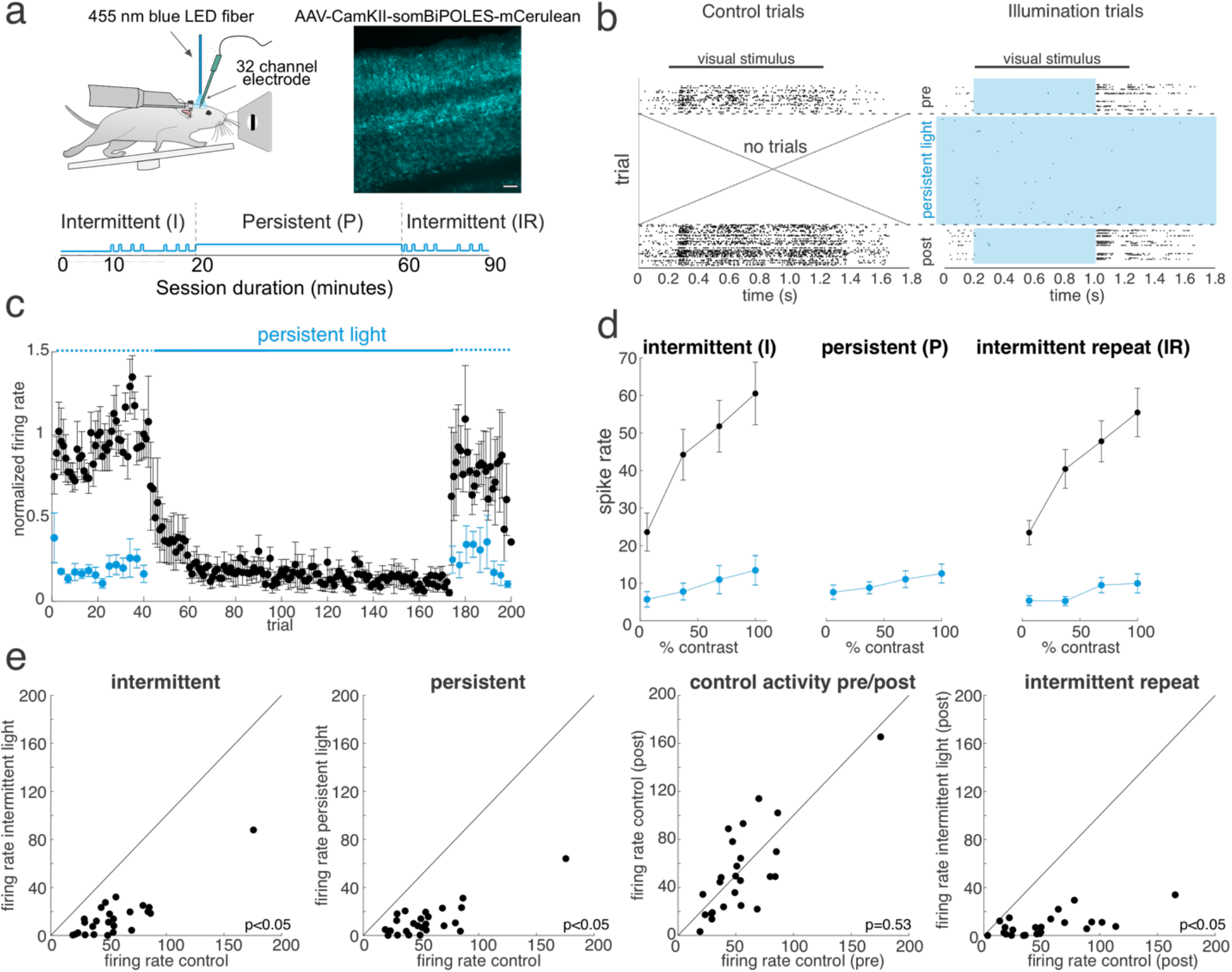
Acute and persistent optogenetic suppression of cortical activity. **A)** Left: schematic of the recording configuration. Right: example image of a post-mortem slice of visual cortex expressing the inhibitory opsin throughout the cortical layers. Scale bar: 100 μm. Bottom: schematic of the illumination procedure with an intermittent illumination block, followed by 40 minutes of continuous persistent illumination, followed by a return to intermittent illumination. **B)** Example raster plot for all spikes on an example electrode during the acute/persistent optogenetic suppression protocol. Left: trials with no illumination during the intermittent phases. Right: trials during the illumination phases (blue rectangle): intermittent (pre), persistent, and intermittent (post), as indicated by the blue shaded regions. **C)** Averaged normalized spiking activity across all recorded tetrodes across all experiments (3 mice, 6 sessions, 25 tetrodes) during each phase of the optogenetic protocol for the 100% contrast stimulus. Intermittent illumination for ∼40 trials, followed by continuous illumination for ∼130 trials, followed by a return to intermittent illumination. Black dots correspond to no light trials; blue dots indicate illumination trials. Each phase of illumination is schematized by the blue bars above the plot; broken lines: intermittent; solid bar: persistent light. **D)** Average contrast response plots across all recorded units in each phase of the silencing experiment. Left: control (black) and intermittent illumination trials (blue). Middle: persistent illumination trials. Right: return to the intermittent illumination. **E)** Scatter plots of recording unit firing rates between the indicated conditions; 3 mice, 6 sessions, 25 tetrodes; Wilcoxon signed rank test. All error bars are s.e.m.

### Acute and persistent effects of V1 inactivation in a visual detection task

Equipped with this flexible optogenetic approach for either intermittent or sustained optogenetic inactivation, we probed how acute versus persistent optogenetic inactivation of V1 would impact visual behavior. We trained somBiPOLES-expressing mice on the same visual contrast detection task used above^16,26^ (Fig. 4a). As before, intermittent silencing of V1, now via excitatory neuron suppression, severely impaired performance (Fig. 4b, Fig. 4d, Fig. S2; p<0.001; n=13 mice). This effect was consistently large: for all 13/13 mice the average (over 6 days) light-on detection threshold was ≥ 1.5x light-off threshold (Table 1, Fig. S2; mean and s.e.m. fold change (c50_I-on_/c50_I-off_): 3.0 ± 0.2). Over several days of intermittent illumination, the reduction of performance was stable, revealing no significant recovery (Pearson r of c50_I-on_ vs day: p=0.52, r=-0.07).

**Figure 4.**
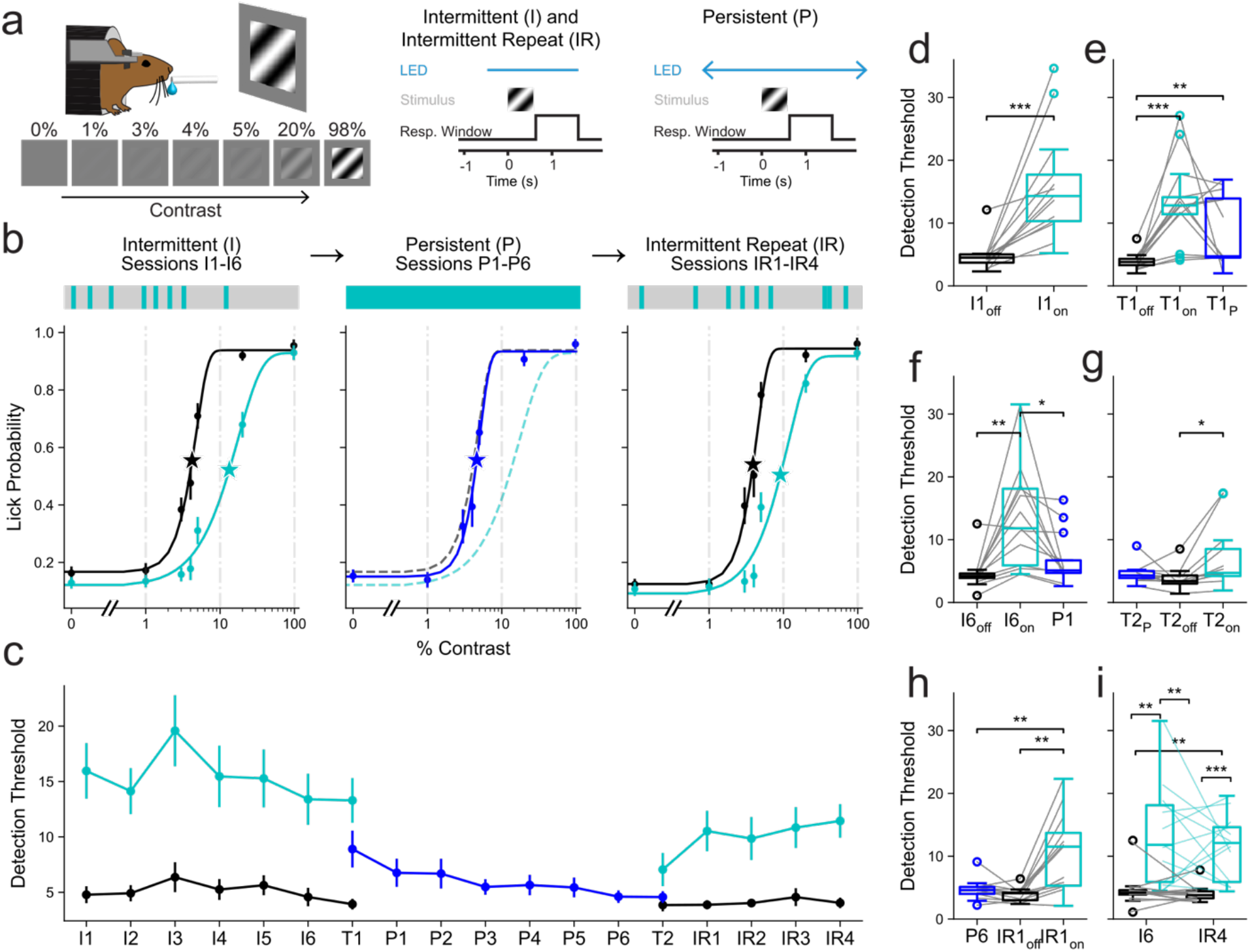
Reversible recovery from persistent inactivation of visual cortex. **A)** Left: Schematic of visual detection task. Right: Optogenetic trial structure: Light is on transiently (∼2 seconds) for 33% of trials (intermittent and intermittent repeat, left) or chronically (persistent, right) for the duration of the session. Gray bar represents stimulus presentation time, black line represents response window, time referenced to the start of the stimulus window. **B)** Top, schematic of silencing blocks, excluding transition days. In the intermittent blocks, silencing was applied on a subset of trials. In the persistent block, silencing was continuous throughout each session. The intermittent repeat block had the same structure as the intermittent block. Bottom, psychometric curves of performance for n = 10 mice tested at the most common contrast set. Stars indicate the fit detection threshold of this average curve. Left, intermittent block, light blue shows light on performance and black is light off performance. Center, persistent block, performance during silencing shown in dark blue, with dashed lines indicating intermittent performance replotted from the left panel for comparison. Right, intermittent repeat block, colors as in intermittent. **C)** Detection threshold on each day of the testing protocol. Transition days (T1 & T2) had one half of the session intermittent, one half persistent. n=13 mice. **D)** Detection threshold in light off (black) vs light on (blue) trials for intermittent day 1 (I1) (p<0.001, n=13 mice, paired t-test). **E)** Detection threshold for T1. In the first half of the session intermittent silencing occurred on light on (light blue) trials interspersed within light off (black) trials. In the second half light was turned on for the entire period for persistent silencing (dark blue). p<0.001, repeated measures ANOVA (rmANOVA). Pairwise comparisons: I_on_ vs I_off_: p<0.001, I_on_ vs P: p=0.08, I_off_ vs P: p<0.01; multiple-comparisons corrected paired t-test; n=13 mice. **F)** Detection threshold during light off vs light on trials for intermittent day 6 (I6) vs persistent day 1 (P1). p<0.001, rmANOVA. Pairwise comparisons: I6_on_ vs P1: p<0.05, I6_on_ vs I6_off_: p<0.01, I6_off_ vs P1: p=0.12, multiple-comparisons corrected paired t-test; n=13 mice. **G)** Detection thresholds for T2, which is the inverse of T1 (e). p<0.05, rmANOVA. IR_on_ vs P: p=0.14, IR_on_ vs IR_off_: p<0.05, IR_off_ vs P: p=0.24; n=13 mice. **H)** Detection thresholds in persistent day 6 (P6) versus intermittent silencing day 1 (IR1). p<0.01, rmANOVA. IR1_on_ vs P6: p<0.01, IR1_on_ vs IR1_off_: p<0.01, IR1_off_ vs P6: p=0.10, multiple-comparisons corrected paired t-test; n=13 mice. **I)** Comparison of detection thresholds for intermittent day 6 (I6) and intermittent repeat day 4 (IR4). p<0.001, rmANOVA. Pairwise comparisons: Ioff vs Ion: p<0.01, Ioff vs IRoff: p=0.48, Ioff vs IRon: p<0.01, Ion vs IRoff: p<0.01, Ion vs IRon: p=0.48, IRoff vs IRon: p<0.001; multiple-comparisons corrected paired t-test; n=13 mice. B-C) Data are shown as group mean ± s.e.m. D-H) Boxplots show the median (center line), interquartile range (box limits: 25th to 75th percentile), and whiskers (extending to the most extreme data points within 1.5× the interquartile range). Outliers beyond the whiskers are shown as individual points.

**Table 1.** Individual mouse performance comparison between conditions, per stage. Each cell gives the ratio of contrast detection thresholds (c50) between two conditions for one mouse, averaged across all sessions of each block: intermittent silencing (I, 6 sessions), persistent silencing (P, 6 sessions), and intermittent repeat (IR, 4 sessions), with light-on and light-off trials denoted by subscripts. Values >1 indicate a higher threshold (worse detection) in the numerator condition; values <1 indicate a lower threshold. Bold entries and saturated colors mark ratios differing at least 1.5-fold from unity in either direction (≥1.5× or ≤0.67×). The bottom row counts the mice meeting that criterion in each comparison.

|  | I<sub>on</sub> / I<sub>off</sub> | P / I<sub>on</sub> | P / I<sub>off</sub> | P / IR<sub>on</sub> | P / IR<sub>off</sub> | IR<sub>on</sub> / IR<sub>off</sub> |
|---|---|---|---|---|---|---|
| mouse 1 | **2.2×** | **0.5×** | 1.0× | **0.4×** | 1.0× | **2.4×** |
| mouse 2 | **2.9×** | **0.3×** | 1.0× | **0.4×** | 1.1× | **2.5×** |
| mouse 3 | **3.5×** | **0.4×** | 1.4× | **0.5×** | 1.3× | **2.9×** |
| mouse 4 | **2.9×** | **0.4×** | 1.1× | **0.5×** | 1.0× | **2.1×** |
| mouse 5 | **3.7×** | **0.6×** | **2.2×** | 1.1× | **2.5×** | **2.2×** |
| mouse 6 | **3.0×** | **0.3×** | 0.8× | **0.5×** | **2.2×** | **4.4×** |
| mouse 7 | **2.2×** | **0.3×** | 0.6× | **0.5×** | 1.1× | **2.1×** |
| mouse 8 | **1.6×** | 0.7× | 1.2× | 0.8× | 1.3× | **1.7×** |
| mouse 9 | **3.6×** | **0.5×** | **1.9×** | **0.6×** | **1.7×** | **2.7×** |
| mouse 10 | **4.6×** | **0.2×** | 0.9× | **0.4×** | 1.3× | **3.1×** |
| mouse 11 | **2.3×** | **0.5×** | 1.2× | **0.3×** | 0.9× | **3.1×** |
| mouse 12 | **3.2×** | **0.3×** | 1.0× | 0.9× | **1.7×** | **1.8×** |
| mouse 13 | **2.7×** | **0.3×** | 0.8× | **0.6×** | 1.3× | **2.1×** |
| **# mice<br>1.5× difference** | **13** | **12** | **2** | **10** | **4** | **13** |

To better mimic a permanent lesion, we then illuminated V1 for the entire duration of the behavioral session over several successive days. Notably, this avoids irreversibly lesioning the tissue so that silencing can be switched off in future sessions. Moreover, it avoids silencing V1 outside the task, such as in the home cage, which might on its own cause local or downstream circuit reorganizations^27^. It may also be preferable to reversible silencing by agents, such as muscimol, as the onset and offset of optogenetic suppression can be precisely controlled, and its spatial extent can be at least partially demarcated by the extent of opsin expression (Fig. S2b).

We applied this persistent illumination scheme to the same mice as above to test if mice might recover performance over time despite continued suppression of V1. Between the last day of intermittent silencing and the first day of persistent silencing, we included a ‘transition day’ in which the first half of the session employed intermittent silencing, and the second half employed persistent silencing to look for fast changes in the role of V1 in behavior. On average, performance during the persistent silencing was not significantly different from light on performance in the intermittent portion of the transition day (Fig. 4e; Transition Day 1 (T1); Intermittent light on (I_on_) vs Persistent: p=0.08, I_on_ vs I_off_: p<0.001, I_off_ vs P: p<0.01; n=13 mice). This shows that recovery was not immediate within a session.

We then examined if mice recovered the next day (P1), a full day of persistent silencing. Beginning on this day, the optogenetic light was on for the entirety of each session for six consecutive days (session duration: 57±13 minutes; mean ± SD). On this second exposure to persistent silencing, detection threshold under persistent silencing was not significantly different than the control light-off detection threshold during intermittent silencing (Fig. 4f-h; Intermittent silencing day 6 (I6) vs persistent day 1 (P1); I6_on_ vs P1: p<0.05, I6_on_ vs I6_off_: p<0.01, I6_off_ vs P1: p=0.12; n=13 mice). While, on average, mice recovered by this second exposure, recovery evolved over several sessions of persistent silencing (Pearson r for persistent c50 versus session number (including T1): −0.34, p<0.001). Persistent recovery was consistent across mice: averaged over the six intermittent and persistent days, 12/13 mice showed intermittent light-on detection thresholds 1.5-fold greater than their persistent thresholds (Table 1; intermittent c50 ≥1.5x persistent c50 in 12/13 mice). Of these 12 mice, 10 recovered to their initial light-off performance (Table 1; persistent c50 < 1.5x light-off c50). These data demonstrate that mice can recover much of their contrast sensitivity even during the continued suppression of V1 activity. Along with the recovery observed after mechanical lesion, this suggests that normal V1 activity is not absolutely required for task performance. Thus, so far, the data leave it unclear whether V1 is instructive or permissive in the task.

### Reversion to a reliance on V1 activity after cessation of persistent silencing

We reasoned that we could address this key question by reverting the persistently silenced and behaviorally recovered mice to intermittent silencing. If intermittent silencing of V1 no longer had a behavioral effect, this would imply that the cortex is probably not the default circuit instructing performance on this visual task and is merely permissive. In this case, V1 activity may only be important for ensuring that the other region that instructs the task has sufficient sensitivity to retinal drive, and in the continued absence of V1 input this other brain area homeostatically compensates. Conversely, if mice instead return to failing on intermittent V1 silencing trials after cessation of persistent silencing, this would support the notion that V1 is the default circuit that actively instructs perceptual behavior.

To discriminate between these possibilities, we reverted the persistently silenced animals to intermittent silencing of the cortex only on 33% of trials, as before (intermittent repeat, IR). To resolve any fast changes in behavioral performance, we started with a second transition day composed of persistent silencing in the first half and intermittent silencing in the second half. During the intermittent half of the second transition session, there was already a significant difference between light off and light on (Fig. 4g; Transition Day 2 (T2); Pairwise comparisons: IR_on_ vs P: p=0.14, IR_on_ vs IR_off_: p<0.05, IR_off_ vs P: p=0.24; n=13 mice). However, IR_on_ was not different than P, indicating that within a day reversion was at most partial. Then we began the intermittent repeat block, consisting of four sessions of intermittent silencing alone. On the first day of the intermittent repeat (IR1) block, on average mice returned to showing a dramatic reduction in performance during intermittent silencing trials (Fig. 4b,c,h; IR1_on_ vs P6: p<0.01, IR1_on_ vs IR1_off_: p<0.01, IR_off_ vs P6: p=0.10; n=13 mice). By the end of the intermittent repeat sessions (IR4), the performance in intermittent light-on trials was the same as at the end of the original intermittent silencing block (I6), before the behavioral recovery during persistent illumination (Fig. 4i; p=0.48, paired t-test, n=13 mice). All 13 mice had worse performance in IR light-on than light-off (Table 1, 1.5x larger c50-IR_on_ than c50-IR_off_; mean/sem: 2.6 ± 0.2). Of the 12 mice that recovered, 10/12 fully reverted to requiring V1, shown by worse performance in IR_on_ than persistent light on (Table 1, 1.5x larger c50-IR_on_ than c50-P). Thus, in the great majority of animals, effects during intermittent repeat and intermittent silencing were substantially larger than during persistent silencing, indicating a reversion to requiring V1 even after learning to perform without it. This reversion to requiring V1 activity is consistent with V1 playing an instructive rather than a merely permissive role in the visual task.

An advantage of the flexible optogenetic paradigm used here is that V1 has normal activity between each behavioral session, which can help further disambiguate between instructive and permissive roles. If homeostatic recovery of other brain areas explains behavioral recovery during V1 silencing, we would expect that recovery would revert between persistent silencing sessions (when V1 activity is normal). To test this, we analyzed performance across persistent silencing sessions (Fig. S3). If homeostatic recovery were reverting while V1 is active and then rapidly re-adapting at the start of sessions, performance should be worse at the start of each persistent silencing session and recover by its end. However, hit rates in the first fifth of a persistent session were comparable to those at the end of the prior session (Fig. S3a, p=0.76, n=13 mice), suggesting that whatever recovery occurs, it does not occur repeatedly within each session. This provides additional evidence against a homeostatic explanation for the behavioral recovery during persistent silencing, implying instead that the mice learn to use an auxiliary brain area in the absence of V1 activity.

### The superior colliculus supports visual detection during persistent V1 silencing

We can consider multiple explanations for why mice could recover their original performance levels despite continued silencing of V1 activity. They could have exploited another brain region also receiving retinal input, or they could have learned to use any residual V1 activity (within the silenced region or at the margins). Given the substantial recovery after lesion in both our and other studies, we opted to first look for a compensatory brain region. Several lines of evidence supported focusing on the superior colliculus (SC): it is the other major target of retinal output, silencing of the SC can lead to visual deficits in mice including in contrast detection^28,29^, and in rats SC can compensate for loss of V1 following a permanent lesion^12^. We reasoned that if the SC compensated for the loss of V1 activity in the recovered state, optogenetic suppression of SC during V1 persistent silencing should cause dramatic reductions in behavioral performance. Such a result would also help rule out alternative explanations for persistent recovery.

To test this notion, we generated a new cohort of animals that allowed separate and simultaneous silencing of SC and V1. We expressed somBiPOLES in V1 and ChR2 in GABAergic inhibitory neurons in the superior colliculus^28,29^ (Fig. 5a, Fig. S4b). In this cohort, we quantified performance using the hit rate rather than c50 because the combined SC + V1 silencing impaired behavior in some mice so much that we could not fit a psychometric curve (Fig. S4a). We first silenced the two regions separately. Silencing SC alone significantly reduced hit rate (Fig. 5b, p<0.01, n=4 mice; hit rate, light off: 67 ± 5%, SC light on: 54 ± 3%), and silencing V1 alone also reduced hit rate, as expected from our previous experiments (Fig. 5c, p<0.01, n=4; hit rate, light off: 67 ± 6%, V1 light on: 45 ± 5%). Neither SC nor V1 silencing entirely abolished task performance, indicating that multiple pathways might normally support contrast detection in intact mice.

**Figure 5.**
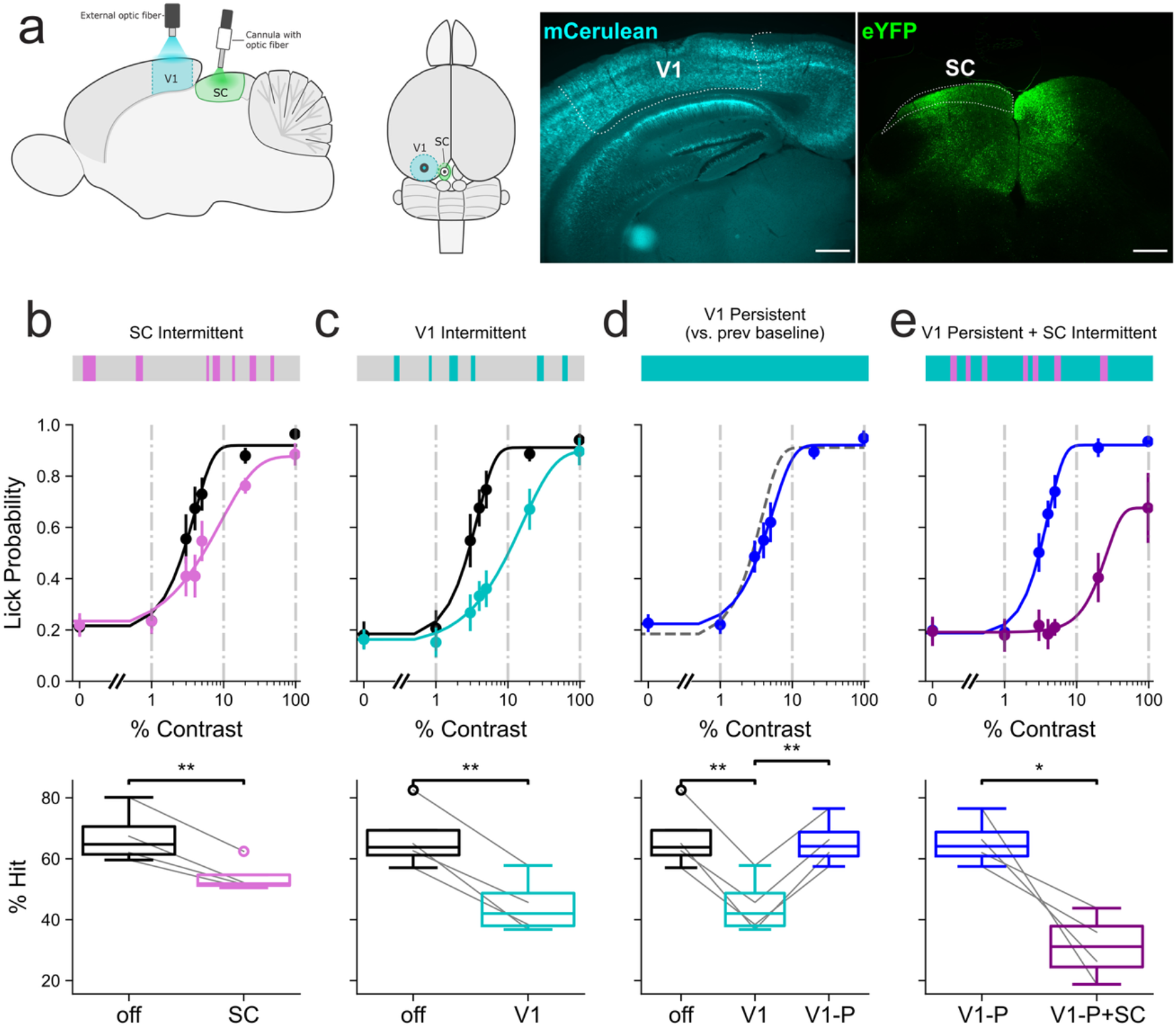
Superior colliculus silencing abolishes recovery from persistent silencing. **A)** Left, schematic sagittal view of the experimental setup, in which V1 and SC can be silenced separately or simultaneously. Middle, dorsal view showing the thinned-skull area with the optic fiber over V1 and the cannula with optic fiber over SC. Right, expression of mCerulean (somBiPOLES) in V1 and eYFP (ChR2) in inhibitory neurons in SC. Scale bar: 500 μm. For additional mice see Fig. S4. **B-E)** Top, schematic of the trial structure for silencing V1 (blue) and SC (purple). Middle, psychometric curves averaged over all mice for sessions in that block (n=4). Data are shown as group mean ± s.e.m. Bottom, comparison of hit rates between experimental conditions. Boxplots show the median (center line), interquartile range (box limits: 25th to 75th percentile), and whiskers (extending to the most extreme data points within 1.5× the interquartile range). Outliers beyond the whiskers are shown as individual points. **B)** Comparison of light off control versus intermittent SC silencing; p<0.01, paired t-test, n=4 mice. **C)** Comparison of light off control versus intermittent V1 silencing; p<0.01, paired t-test, n=4 mice. **D)** Comparison of light off control (off) and intermittent light on (V1) from (C) versus persistent V1 silencing (V1-P); p<0.001, rmANOVA. Pairwise comparisons: V1 vs off: p<0.01, V1 vs V1-P: p<0.01, V1-P vs off: p=0.60; multiple-comparisons corrected paired t-test; n=4 mice. **E)** Comparison of V1 persistent (V1-P) and V1 persistent with intermittent SC silencing in the same session (V1-P+SC). p<0.05, paired t-test, n=4 mice.

Next, we tested the hypothesis that SC compensates for V1 when it is inactivated in persistent silencing by silencing SC intermittently during persistent V1 silencing. We persistently silenced V1 for several days, as before, and the mice recovered their performance, as expected (Fig. 5d, pairwise comparisons: V1-intermittent (V1-I) vs V1-Persistent (V1-P): p<0.01, V1-P vs off: p=0.60, n=4; hit rate, V1-P: 62 ± 4%). If task performance in this recovery period depends on the SC, silencing it on top of continued persistent V1 silencing should reveal a pronounced deficit. Indeed, when we suppressed SC concurrently with persistent V1 silencing, we observed a profound reduction in performance (Fig. 5e, p<0.05, n=4; hit rate, V1-Persistent: 66 ± 4%, V1-Persistent + SC: 31 ± 5%). This argues that much of the recovery in these mice during persistent silencing of V1 can be attributed to compensatory action of the SC. The behavioral deficit when silencing SC during V1 persistent silencing trended towards being larger than silencing SC alone (p=0.058, n=4 mice; hit rate, SC silencing alone: 54 ± 3%, SC silencing during V1-Persistent: 31 ± 5%), suggesting that mice are more reliant on SC when V1 is also absent. Perhaps just as importantly, these results make alternative explanations for the behavioral recovery under persistent V1 silencing less likely: for example, if the mice had learned to exploit residual V1 activity to recover their task performance, or had learned to move their eyes to exploit a non-silenced region of the visual cortex, silencing the SC should likely not have had such a strong, if any, effect. Thus, we conclude that mice recover performance during persistent optogenetic silencing of V1 by instead exploiting activity in SC.

## DISCUSSION

Here, we exploit a simple and potentially broadly applicable three-stage paradigm for using optogenetics to help disambiguate whether a given brain circuit is instructive or permissive in a behavior. In stage one, acute inactivation assesses whether the target circuit is involved at all but does not discriminate between an instructive or permissive role. In stage two, sustained inactivation assesses whether the behavior can recover, either due to relearning with alternative circuits or due to homeostasis. Finally, when there is behavioral recovery, stage three returns to acute inactivation to help determine whether the targeted circuit is instructive or permissive. Additionally, direct simultaneous optogenetic targeting of a putative area responsible for recovery during persistent silencing of the first can more unambiguously resolve the sites of instructive and permissive action in the brain for a given behavior.

Our data on mouse V1 supports a model in which V1 is instructive in perceptual behavior and not merely permissive. The result from the return to intermittent inactivation best distinguishes between the multiple competing explanations for why acute V1 suppression impairs behavior, but persistent silencing yields recovery. First, the performance recovery could have been explained by homeostatic recovery of a downstream area (e.g., the SC) whose normal activity level depends on sensory cortical input. If this were true, then when the mice were reverted to intermittent silencing, one would have expected the mice to be resistant to cortical inactivation since the downstream area had homeostatically become independent of cortical drive. However, this was not the case, as mice quickly reverted to depending on V1 activity. It is also possible that the putative homeostatic recovery could itself have disappeared in the intervening time between the last persistent inactivation session and the first intermittent inactivation session of the subsequent block. However, if this were true, it should have happened between each of the persistent inactivation sessions, and then mice should have shown transient poor performance at the beginning of each persistent inactivation session. But this was also not the case (Fig. S3a), arguing against any spontaneous recovery between days of persistent illumination. Additionally, our data put constraints on the timescale of recovery during persistent inactivation. Our results show the recovery was not fast but developed progressively over several days. This is not consistent with a fast homeostatic mechanism that might rebalance activity on short timescales. Thus, on the whole, homeostatic recovery is not consistent with our results.

In a second scenario, the recovery of performance during persistent silencing could have been explained by the animals relearning the task using alternative circuits that do not depend on primary sensory cortical activity^5^. In this scenario, task-relevant information is available to the animal in both the primary sensory cortex and alternative structures independently, and though originally the animal depended on V1 activity to solve the task, over several days of training in its absence, the animal relearned the task via these alternate circuits. This was supported by the recovery over several sessions during persistent silencing.

Our data further argue that V1 is also a favored or default pathway for visual perceptual information because this learned reliance on alternate circuits is impermanent: when primary sensory cortical activity again becomes available, the downstream circuits revert to ignoring this alternative source of information even on trials in which the primary sensory cortex is acutely inactivated. Our results support this scenario; they argue that V1 is the default source of instructive information for solving a simple contrast detection task, while the SC is an accessory visual region that can support low contrast detection in the absence of V1. Tellingly, when persistent silencing of V1 is switched off, the animals return to relying on it. This argues that the adaptive reliance on SC during persistent V1 silencing does not become the default, suggesting that the brain flexibly chooses its source of instructive input, but in this task, it favors V1. Why the mouse brain reverts to using V1, even when SC could support nearly indistinguishable performance, is not yet clear. V1 might generally be a more powerful or reliable source of retinal information to perceptual decision circuits, and when available, these decision circuits will rely on it despite alternative sources of information.

A third possibility is that mice adapted to use residual V1 activity, either through circuit mechanisms or through behavioral strategies such as moving their eyes to avoid the optogenetic scotoma (i.e., the retinotopic region of V1 that was inactivated). Several features of mouse behavior and aspects of the data argue against this. First, saccades in head-restrained mice are small compared to the size of the stimulus (18°) we used, making a gaze-based strategy an implausible route to recovery^30^. Second, post-mortem histology showed broad somBiPOLES expression covering all or nearly all of V1, and our illumination system likely inactivated much, if not all, of V1 based on similar approaches^31^. Thus, it would be surprising if incomplete optogenetic silencing left a sufficiently large and active spared portion of V1 that was within retinotopic reach of our small stimulus consistently across all 12 of 13 animals that recovered from persistent silencing. Third, and most importantly, if in the optogenetic condition the mice had used a strategy that still relied on V1, then the persistent V1 silencing plus SC silencing should have had no effect, but it profoundly impaired mouse performance. While we cannot rule out that some mice may have used residual V1 activity as part of their recovery, the data strongly supports that other pathways (i.e., the SC) are involved.

How should we understand the recovery from permanent lesions and why was recovery only partial? Our new optogenetic results may help us interpret the lesion studies. Our data imply that mice recover from V1 lesion because they can learn to exploit the remaining visual pathway through the SC, at least for the stimuli (i.e., the spatial frequencies) we used. Under persistent silencing, recovery was nearly complete while after lesion it was partial, perhaps because LGN degenerates after aspiration of V1 but not after persistent optogenetic silencing. Interestingly, recovery after lesion was incomplete in this study despite using relatively lower spatial frequencies, which may not have been expected to be as V1-dependent based on prior work^14^. This may reflect differences in stimulus size or in training mice at near-threshold contrasts, where residual deficits are more readily detected, or in LGN degeneration after lesion. Future work employing this approach across a broad set of spatial frequencies, other types of stimuli, and in discrimination tasks, could yield additional insight into the conditions and contexts in which V1 is essential for task performance, or can be partially or completely compensated for by pathways through the SC.

More generally, this multi-stage optogenetic approach can help address the challenges posed by potential off-target effects of acute circuit perturbations and by confounds of permanent lesions. In principle, any study already using optogenetic inactivation can execute this paradigm without substantial additional technical complexity, although it is important to confirm the efficacy of sustained optogenetic suppression as many optogenetic actuators may not be suitable. The main advantage of the three-stage silencing approach used here is that it should lead to more interpretable results and greater flexibility in silencing brain regions during certain stages of a paradigm, such as during learning, to illuminate the interplay between circuits. Moreover, it may aid in understanding the mechanisms of behavioral recovery after brain lesions^27^, helping to delineate the default versus accessory circuits that can contribute to specific behaviors.

## ACKNOWLEDGEMENTS

The authors thank J. Beyer and K. Gopakumar for technical support, N. Bhatla for performing V1 suction lesions, and K. Liang for histology support. This work was funded by NIH R01EY023756 and RF1MH120680. H. Adesnik is a Chan Zuckerberg Biohub investigator. J.S.W. was funded by the German Research Foundation (SPP 1926, Project Nr.: 315380903 WI4485/3-2) and acknowledges Kathrin Sauter and Ingke Braren for technical support.

## AUTHOR CONTRIBUTIONS

D.Q., J.B., H.A.B. and H.A. conceived the experiments. D.Q. performed all surgical procedures and behavioral experiments with assistance from M.W. H.A. performed all electrophysiological recordings. H.A.B. developed the task and performed quantitative analysis. J.B. performed an initial series of behavioral and optogenetics experiments to define illumination parameters. J.S.W. developed and contributed the AAV-somBiPOLES vector. H.A. wrote the manuscript with help from D.Q. and H.A.B.

---

**Figure S1.**
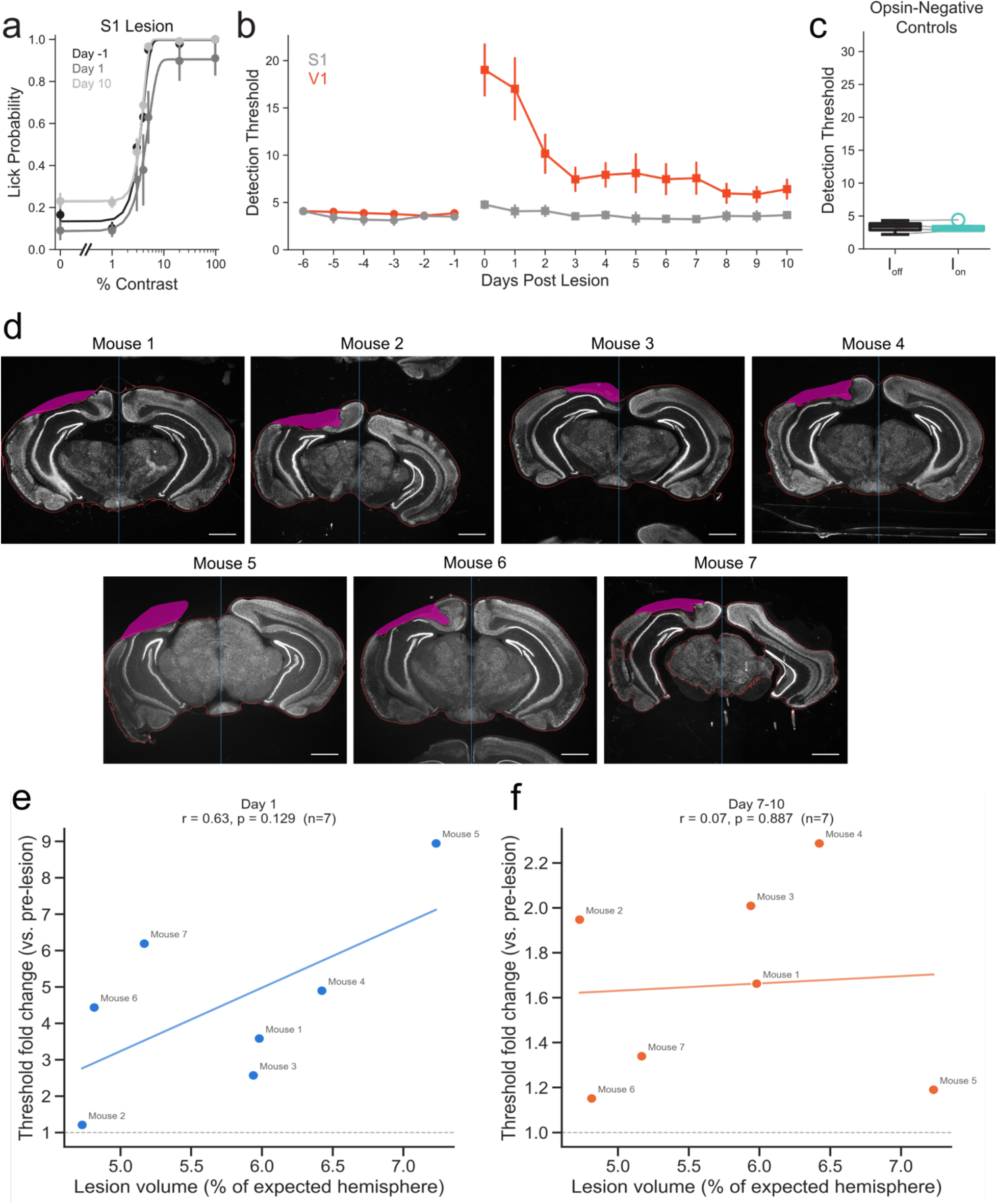
Control experiments for the specificity of V1 lesion and optogenetic inactivation on contrast detection, and visualization of lesion extent. **A)** Psychometric curves of performance of mice with S1 lesions on the day immediately prior to lesion (Day −1, black), day after lesion (Day 1, dark gray), and Day 10 (light gray) after lesion. Lines, Weibull function fits of the psychometric curve. **B)** Performance of mice for six days before lesion (circles), tested hours after lesion (Day 0), and then tested for ten days after lesion (squares). Orange, V1 lesion, n=7 mice; gray, S1 lesion, n=3 mice. **C)** Boxplot showing detection thresholds in opsin-negative controls averaged over 3 sessions per mouse (p = 0.85, paired t-test, n=4 mice). **D)** NeuN-immunostained coronal sections from each of 7 mice displaying the manually traced lesion (magenta), segmented hemisphere boundary (red), and manually identified hemisphere midline (blue) used to compute lesion volume as percent of expected hemisphere. Scale bar: 1 mm. E) Lesion volume vs. detection-threshold fold change (day/pre, linear ratio) at Day 1, r=0.63, p=0.129, Pearson correlation, n=7 mice. F) Lesion volume vs. detection-threshold fold change at Day 7–10, r=0.07, p=0.887, Pearson correlation, n=7 mice.

**Figure S2.**
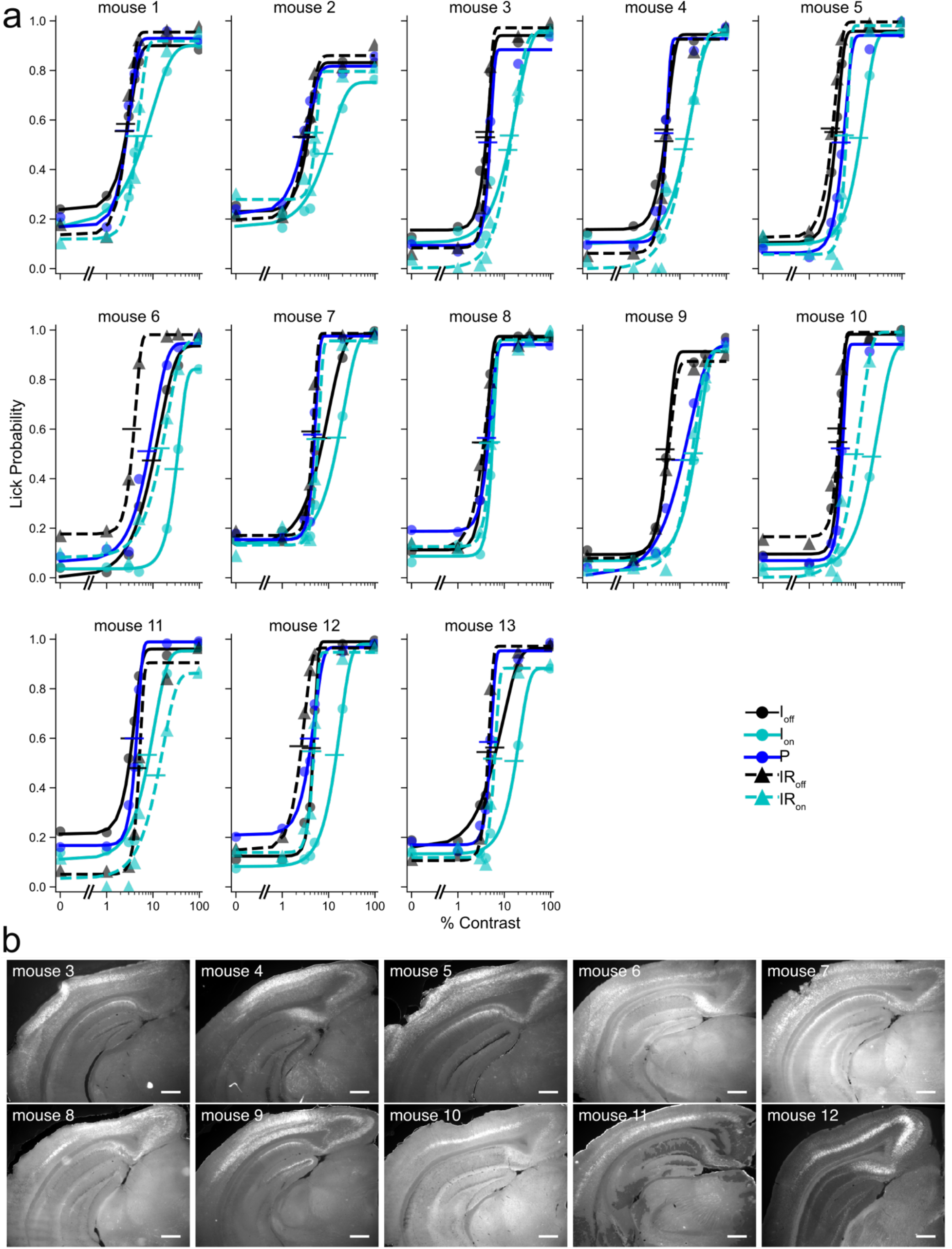
Behavioral performance curves and histology for each mouse. **A)** Psychometric curves of performance for each mouse during each stage of the task, fit with a Weibull function. Data are group mean. Horizontal bars indicate detection threshold/c50 (arbitrary width). **B)** Images showing mCerulean somBiPOLES expression in V1 for 10 of the 13 mice in A for which histology images were available. Scale bar: 500 μm.

**Figure S3.**
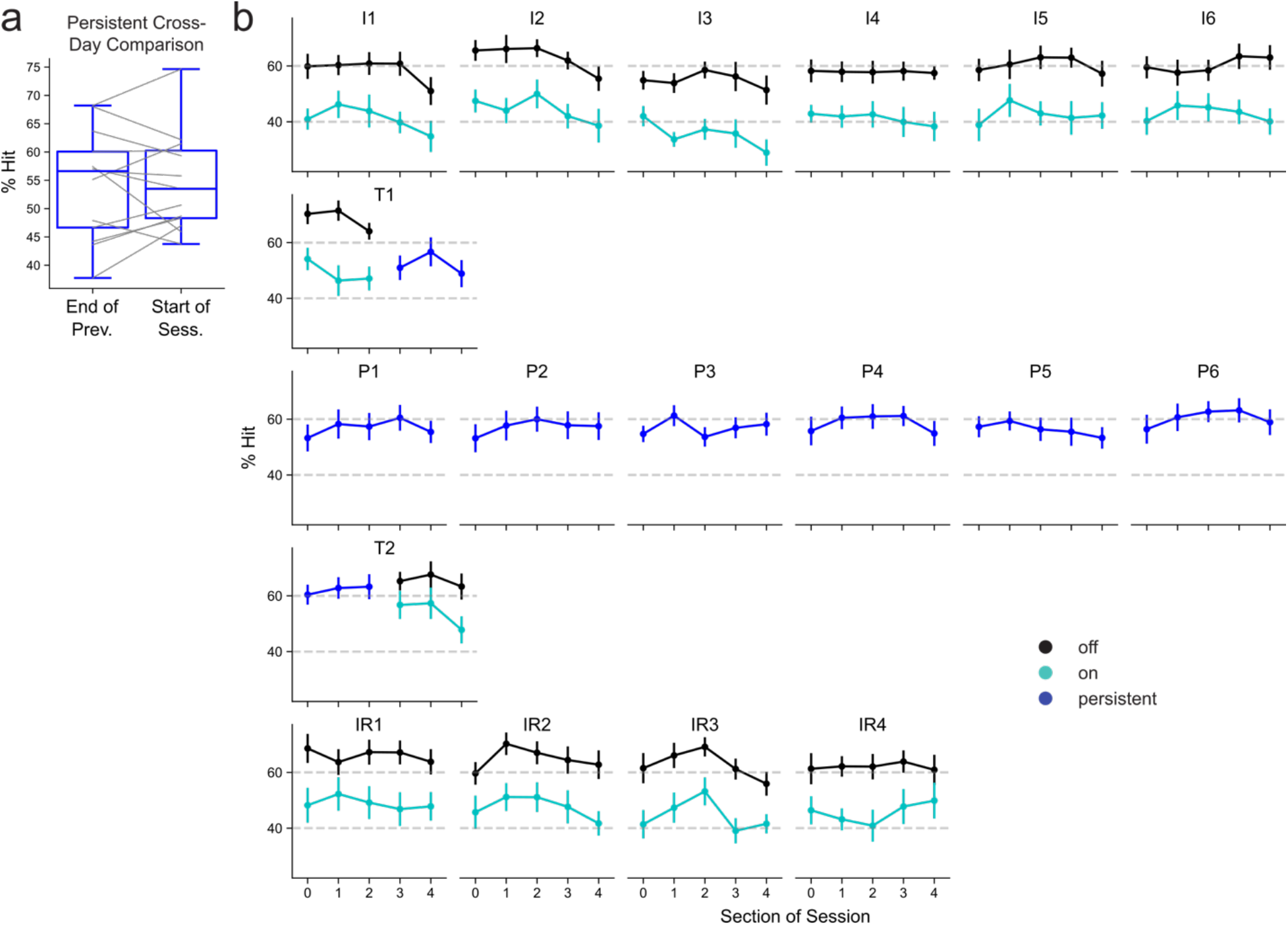
Within-session performance comparisons. **A)** Comparison of, for persistent silencing days, last bin of performance on prior days vs bin on next day; p=0.76, paired t-test; n=13 mice. **B)** Binned hit rate (rate of response for all trials with contrast >0%) for sections of sessions. Non-transition days were split into five sections, transition days into three sections per sub-session. From top to bottom: Intermittent sessions 1-6, transition day 1, persistent sessions 1-6, transition day 2, intermittent repeat sessions 1-4. For this hit rate visualization only n=10 mice which were always run with the same contrasts in all stages are shown. Group mean ± s.e.m.

**Figure S4.**
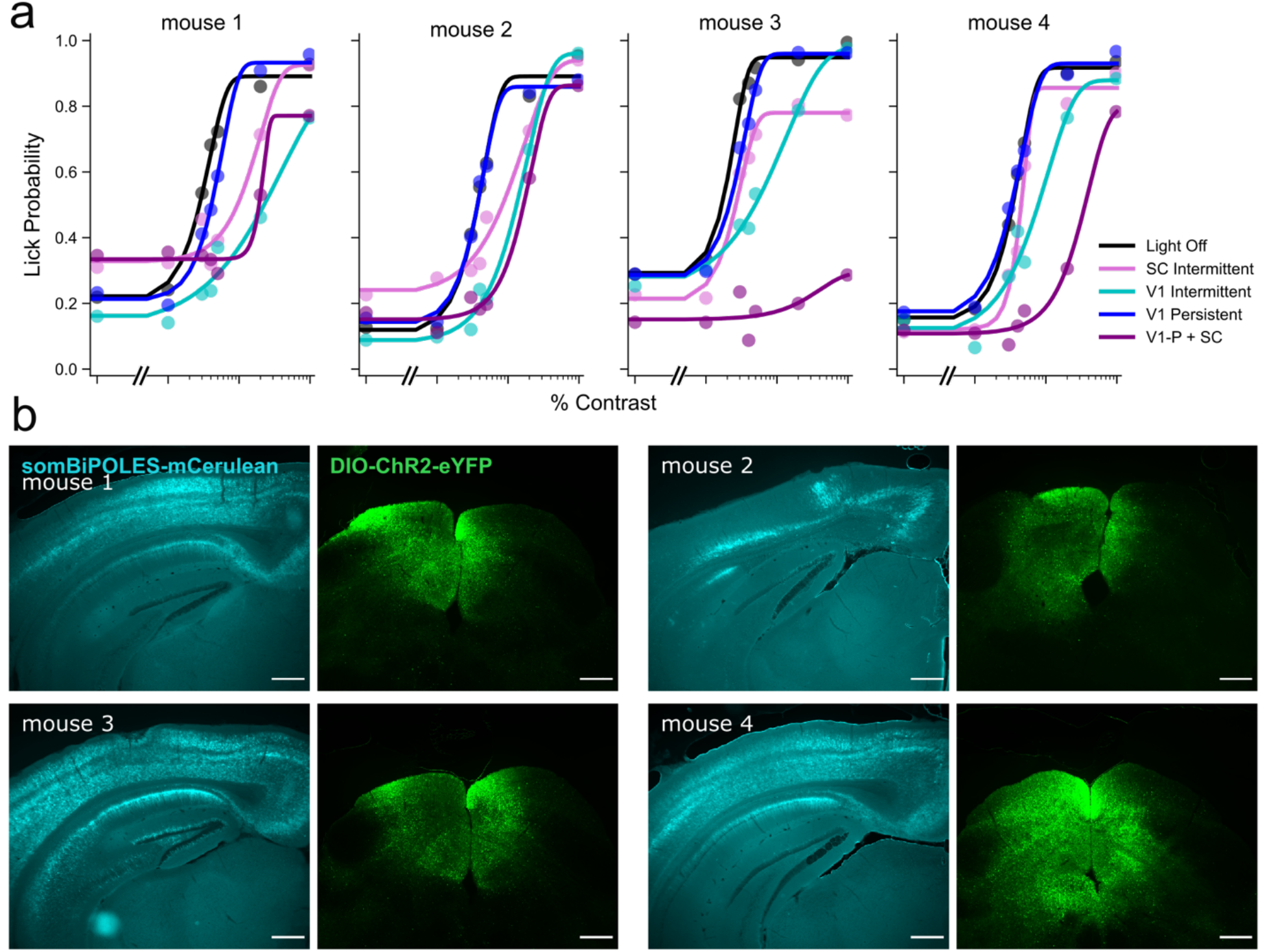
Individual mouse performance and histology for superior colliculus and V1 experiments. A) Psychometric curves for each mouse at different stages of the protocol in Figure 5. For viewing, light off performance is combined between light off for SC Intermittent and V1 intermittent, and V1-P is combined between V1 Persistent alone sessions and V1-P only trials of sessions that included V1 Persistent and SC Intermittent. Data are mean over sessions with fit psychometric curves. B) Histology images for each of the mice in A, showing expression in Gad2-IRES-Cre mice of somBiPOLES-mCerulean in V1 and DIO-ChR2-eYFP in inhibitory neurons in SC. Scale bar: 500 μm.

## MATERIALS AND METHODS

### Animals

Mice used for experiments in this study were C57BL/6J (Jax stock: 000664), PV-IRES-Cre (Jax stock: 008069) crossed with Rosa-LSL-ChR2 (PV-Cre;Rosa-LSL-ChR2(Ai32)) (Jax stock: 012569) and Gad2-IRES-Cre (Jax stock: 010802)^32–34^. Mice were housed in cohorts of five or fewer in a reverse light:dark cycle of 12:12 hours, with experiments occurring during the dark phase. Both female and male mice were used. For optogenetic silencing with somBiPOLES, a total of 13 C57BL/6 mice were used (7 males) aged 10 – 20 weeks. For the V1 silencing in Figure 2, 10 PV-Cre; Ai32 mice were used (5 males) aged 8 – 15 weeks. For lesion experiments and controls, a total of 14 C57BL/6 mice were used (5 males) aged 12 – 30 weeks. For dual V1 and SC experiments, a total of 4 Gad2-IRES-Cre mice were used (2 males) aged 13 – 16 weeks.

### Viral injection

Neonatal C57BL/6 and Gad2-IRES-Cre mice (P3–P5) were cryoanesthetized and their heads were placed in a ceramic mold and secured with non-abrasive tape. Viral solutions (AAV9-CaMKII-somBiPOLES-mCerulean; titer: 1×10¹³ vg/mL; Addgene: 154948-AAV9)^22^ were loaded into beveled glass injection needles (WPI) and mounted in a Nanoject II (Drummond Scientific). Using a micromanipulator (Sutter MP-285), the needle was inserted through the skin and skull over the left primary visual cortex (V1; 2.3 mm lateral and 0.1 mm anterior to lambda) at three locations spaced approximately 0.3 mm apart. At each location, virus was delivered at four cortical depths (400, 300, 200, and 100 μm below the surface), with 20 nL injected per depth at a rate of 10 nL/s.

### Headpost surgery

All experiments were performed in accordance with the guidelines and regulations of the Animal Care and Use Committee of the University of California, Berkeley. For head fixation during behavioral and physiological experiments, a small custom stainless-steel headplate was surgically implanted. Briefly, adult mice (P70 – P155) were anesthetized with 2-3% isoflurane and mounted in a stereotaxic apparatus. Buprenorphine (0.05-0.1 mg/kg SC) was administered for analgesia and dexamethasone (2 mg/kg SC) to prevent cerebral edema. Body temperature was monitored and maintained at 37°C. The scalp was removed, the fascia retracted, and the skull lightly scored with a drill bit. Vetbond (3M) was applied to the skull surface, and the headplate was fixed to the skull with Metabond (C&B). A fine-point marker was used to note the approximate location of the left primary visual cortex (V1; 2.7mm lateral, 0 mm posterior to lambda). For all behavioral experiments involving mice injected with somBiPOLES, a glass cranial window was implanted over the marked site to provide optical access to the cortex. In V1 silencing comparison experiments, the skull was instead thinned and covered with a single layer of Vetbond and another of nail polish to enhance light penetration^35^. Similarly, for electrophysiological recordings, the skull was thinned to transparency on the day of the experiment to ensure optimal electrode access and signal quality. Mice received buprenorphine and meloxicam for pain management and could recover for at least three days before being placed on water restriction.

### SC Viral Injection

To silence the superior colliculus (SC), Gad2-IRES-Cre mice (P50–P75) that had previously received neonatal V1 injections of somBiPOLES underwent a second stereotaxic injection of AAV9-EF1a-DIO-hChR2-eYFP (titer: 2×10¹² vg/mL; Addgene: 20298-AAV9)^36^. The viral solution was loaded into a beveled glass injection needle and mounted in a Micro4 microinjector (World Precision Instruments). Mice were anesthetized following the procedure described above, and a small craniotomy was made over the SC (0.3 mm anterior and 0.5 mm lateral to lambda). The needle was then advanced to two depths (1.4 and 1.2 mm), with 60 nL delivered at each site at a rate of 20 nL/min. Following each injection, the needle was held in place for 10 minutes prior to retraction to minimize viral backflow. The skull was cleaned with alcohol swabs, and the incision was closed with absorbable sutures.

### SC Cannula Implant and V1 Skull Clearing

One week following SC viral injection, a fiber optic cannula (1.25 mm OD, 200 μm core, 0.39 NA, 2 mm length, Thorlabs Inc.: CFMLC12L02) was implanted through the same craniotomy. After induction of anesthesia, the head was positioned at a 15° angle from the midline, and the cannula was advanced to a final depth of 0.9 mm^29^. The cannula was secured with Vetbond (3M) and Metabond (C&B), with care taken to leave the medial aspect unsealed to preserve ipsilateral V1 optical access. Following cannula implantation, a headpost was fixed using the protocol described above. Finally, the skull overlying V1 ipsilateral to the implanted fiber was thinned and cleared for optical access similarly to the comparison experiments method described above.

### Lesion surgery

Water-restricted adult mice (P75-P180) previously headplated (see headpost surgery for procedure) and trained to perform the behavioral paradigm were anesthetized using isoflurane 2.5% and mounted in a stereotaxic apparatus. Buprenorphine (0.05-0.1 mg/kg SC) was administered at anesthetic induction. The eyes of the animal were covered with ocular lubricant to keep eyes moist. A dental burr was used to perform a 3 mm × 4 mm rectangular craniotomy over the left visual cortex (V1; 2.7mm lateral, 0mm posterior to lambda). Exposed cortical tissue was aspirated using a vacuum-attached blunt needle (16-30 awg). The lesion cavity was extended ventrally until the alveus surface of the hippocampus is exposed. The cavity was covered and sealed using a glass coverslip. Animals received meloxicam and buprenorphine post-surgery for analgesia. Mice performed on the behavioral paradigm 2-4 hours post-surgery and performance is reported as lesion day (Day 0).

### Lesion Quantification Analysis

The relationship between lesion size and behavioral decline was assessed by comparing detection threshold change at Day 1 and Day 7–10 post-lesion (see Methods above) with lesion volume. Lesion boundaries were manually traced on NeuN-immunostained coronal sections for each mouse. For each traced section, the hemisphere midline was manually identified, separating the lesioned hemisphere from its intact contralateral counterpart. Whole-hemisphere tissue extent was obtained by Gaussian-smoothed intensity thresholding of the NeuN signal using this boundary as reference for determining the total brain’s slice surface area. Lesion volume was expressed as the summed lesion area across all traced sections divided by the summed hemisphere area (Cavalieri principle), given as a percentage^37,38^. The relationship between lesion volume and threshold change was assessed using Pearson correlation, computed separately for Day 1 and Day 7–10.

### In vivo electrophysiology recordings

Mice were anesthetized with isoflurane (2.5%), and a small craniotomy (<250 μm) was opened over V1 with a fine needle. Mice were transferred to the electrophysiology rig where they were head fixed and allowed to wake and run freely on a circular treadmill. A 32-channel multielectrode array (NeuroNexus, A1×32-5mm-25-177-a32) was guided into the brain using a micromanipulator (Sutter Instruments) and a stereo microscope (Leica). Electrical activity was amplified and digitized at 30kHz (SpikeGadgets Inc.) and stored on a computer hard drive. The bottom of the electrode was zeroed with the surface of the brain to determine z depth. The electrode array was lowered ∼650-750 microns into V1. A multimode fiber (400 μm diameter) was positioned ∼2-3mm from the brain next to the recording electrode to deliver 455 nm LED light during optogenetic illumination at a total of 2mW/mm^2^ measured at the tip of the fiber.

Spike detection was performed across assigned tetrodes (four adjacent electrode contacts) using the SpikeGadgets threshold detection algorithm. Analysis was performed on all detected spikes on each tetrode. Not all assigned tetrodes were used because the top tetrodes were often located in L1 or outside of the brain. Statistically significant differences between conditions were determined using the Wilcoxon signed rank test across all tetrodes. Since mCerulean was not readily visible during adult surgeries, all experiments were initially performed blind to successful opsin expression, and animals with no obvious physiological optogenetic effects (strong activation by red light via the Chrimson in somBiPOLES) were excluded, which correlated with weak or no expression observed post-mortem.

### Water restriction

Initial animal weight was recorded for 3 days before water restriction to establish baseline weight. Mice were then placed on water restriction. On training days, mice received most water during the task. Mice were weighed after training and given additional water if their weight dropped below 70% of initial weight. Food was available *ad libitum*. The weight and health (quality of the fur and nails, gait, posture) of the mice were monitored daily.

### Behavioral training on the contrast detection task

#### Apparatus

For the contrast detection task, the visual behavioral apparatus was controlled in real time using an Arduino DUE (A000067) synchronized with Raspberry Pi 3+/4+ (BCM2837B0/SC0193(9)) and custom software in Python, Arduino and Java^39^. The mouse was head fixed bilaterally and placed on a 3D-printed holder tube. Mouse licking was measured through the detection of changes in the capacitive load in a touch sensor (Adafruit, AT42QT1070) caused by the contact of the mouse’s tongue and a 0.05-inch diameter steel lickport. Water was delivered through the lickport using a 2-way normally closed isolation valve control (Neptune Research Inc.). Visual stimuli were displayed on a gamma-corrected LCD monitor (Podofo, 15.7 x 8.9 cm, 1024×600 pixels, 60 Hz refresh rate).

#### Behavioral training and task design

Head-fixed mice were trained to lick when they detected a visual stimulus of variable contrasts displayed on the monitor. The training started after weight dropped below 80% of the initial weight from water restriction. Prior to training, mice were habituated to head-fixation for two daily sessions lasting 30 minutes each. The behavioral training protocol consisted of four stages.

In stage 1, mice were classically conditioned to lick in response to a visual stimulus that consisted of a square patch with a drifting sinusoidal grating (2Hz, 0.08 cycles/degree, 600 ms, 100% contrast, 34 visual degrees, black background luminance). Each visual stimulus was paired with a water reward delivered at the beginning of a 1000 ms response window. Mice that achieved a performance of more than 90% of correct trials were transitioned to the next phase.

In stage 2, mice were operantly conditioned to respond every time the same stimulus from stage 1 was displayed (2Hz, 0.08 cycles/degree, 600 ms, 100% contrast, 34 visual degrees, black background luminance). Mice received a water reward when they licked within a 1000 ms response window after the visual stimulus presentation. A no-lick period was enforced after two seconds of the trial. Catch trials were introduced and consisted of 25% of total trials. Mice licking in a no-lick period or a catch trial received a 5- to 9-second time-out punishment. Mice achieving performance with a hit rate >90% and false alarm rate <30% for two consecutive sessions were switched to the next phase.

In stage 3, mice continued to train in the same behavioral paradigm with increased background luminance at different sessions until it reached mean luminance. Then, the visual stimulus size was decreased in different sessions until it reached 18 VD. The same performance criteria as in stage 2 were used in every session before transitioning to any change.

In stage 4, mice started to detect stimuli of seven different contrasts, pseudo-randomly distributed. The intensities were customized for each mouse to provide a reliable detection threshold. For PV silencing experiments, contrasts were customized with several variations. For all other experiments, all mice had contrasts 0%, 1%, 3%, 5%, 20% and 98%; a seventh contrast of either 35% or 4% was presented depending on the individual mouse. Mice were introduced to the optogenetic protocol once performance was stable for more than 3 sessions meaning hit rate was above 60-70% with false alarms below 30%. During photostimulation experiments, catch trials were 25% of all trials and the response window was reduced to 500 ms. On average, mice completed 375 trials per session on non-transition days, and 420 trials for the entire session on transition days.

### Optogenetic stimulation during behavior

For V1 one-photon optogenetic experiments, an optical fiber (400 μm diameter, 0.39 NA; Thorlabs Inc.: M98L01) coupled to a mounted 455 nm LED source (Thorlabs Inc.: M455L3) and LED driver (Thorlabs Inc.: LEDD1B) was positioned above the thinned skull or cranial window using a micromanipulator (Narishige: MM3). For SC optogenetic experiments, the implanted cannula was coupled to a patch fiber (200 μm diameter, 0.39 NA; Thorlabs Inc.: M89L01) and a fiber-coupled 470 nm LED (Thorlabs Inc.: M470F4) and LED driver (Thorlabs Inc.: LEDD1B) system via a quick-release interconnect (Thorlabs Inc.: ADAL3). Light output was calibrated daily by measuring power at the fiber tip using a power meter (Thorlabs Inc.: PM160T). Photostimulation power was set to 2.0 mW for somBiPOLES-expressing mice (V1), 4.0 mW for ChR2-expressing mice (SC), 5.0 mW for non-opsin controls, and 2.0–8.0 mW for PV-Cre; Ai32 mice. During the intermittent blocks, optogenetic stimulation was delivered on 33% of all trials, pseudo-randomly distributed in blocks. Light was delivered at a random time prior to the stimulus onset, between 300 - 3000 ms, to avoid cueing the mouse to the trial start. Light was sustained until the end of the response window. During the persistent block, optogenetic stimulation was turned on prior to the beginning of the session and remained on until the session ended. All mice injected only with somBiPOLES on V1 underwent 18 consecutive days of testing consisting of 6, 6, and 4 sessions in the I, P, and IR blocks, respectively, with two additional transition days (T1 and T2) between different blocks. A custom-designed removable light-blocking system was attached to the head of the mouse to prevent the mice from seeing the optogenetic stimulation light. A blue masking light and a plastic eye patch over the left eye were also placed to prevent additional extraneous cues.

For the superior colliculus experiments in Figure 5, mice underwent a modified protocol. First, there were three sessions of intermittent superior colliculus silencing, then three sessions of intermittent V1 silencing, then five sessions of V1 persistent silencing, then three sessions of V1 persistent silencing plus SC intermittent silencing. No transition days were used. Intermittent silencing occurred on 33% of trials as before with the same timing rules.

### Histology

Mice were anesthetized with 5% isoflurane and ketamine in preparation for transcardial perfusion with phosphate-buffered saline (PBS), followed by 4% paraformaldehyde (PFA). Brains were removed, post-fixed overnight in PFA at 4°C, rinsed three times in PBS, then cryoprotected in 30% sucrose for 1–2 days. Brains were monitored to ensure they initially floated and subsequently sank, indicating complete cryoprotection. Afterward, a sliding microtome (American Optical Company), frozen using dry ice and ethanol to ensure proper adhesion and sectioning, was used to obtain 40 μm coronal sections of S1 and/or V1.

For NeuN staining, sections were incubated for 1 hour at 4°C on a rocker in blocking solution containing 3% normal goat serum (NGS), 0.6% Triton X-100, 0.2% Tween-20, and 3% bovine serum albumin (BSA) in PBS. Primary antibody incubation was performed overnight at 4°C using Rabbit anti-NeuN (Abcam) diluted 1:10,000 in blocking buffer. The following day, sections were washed in PBS with 0.25% Triton X-100 (PBS-T) and incubated for 1 hour at room temperature with Alexa Fluor 405–conjugated Goat anti-Rabbit IgG H&L secondary antibody (1:1000; Thermo Fisher Scientific). Sections were washed again in PBS-T, mounted onto glass microscope slides, and allowed to dry before applying coverslips and Vectashield mounting medium (Vector Laboratories) to reduce movement artifacts.

Imaging was conducted using an Olympus MVX10 Stereo Microscope with a 2× and 0.63× objective lens, 1.8× zoom, 0.5 numerical aperture, and 500 ms exposure. Cell counting was performed using a custom QuPath v0.2.3 pipeline automated for detection of NeuN-positive cells within manually defined regions of interest across sequential brain slices, using intensity thresholds and cell segmentation to quantify labeled nuclei.

### Quantification and Statistical Analysis

Hit rate (% Hit) was defined as the number of go trials (contrast > 0) where the mouse correctly licked within the response window divided by the total number of go trials. Days where mice were run on the incorrect contrast levels due to experimenter error were excluded from analysis of hit rate, but not detection threshold. For detection threshold analysis, data for individual sessions were fit using a Weibull cumulative distribution function that fit the false alarm (lower asymptote), lapse rate (upper asymptote), slope and threshold parameter. Functions were fit separately for photostimulation and no-photostimulation trials. Detection threshold was defined as the contrast at which the fit hit rate was halfway between the fit false alarm (lower asymptote) and fit lapse rate (upper asymptote).

Statistical analyses were performed using MATLAB or Python. The analyses performed were repeated measures analysis of variance (rmANOVAs) with spherical correction when sphericity was violated, paired t-tests with Benjamini–Hochberg false discovery rate corrections where needed, independent t-tests, and Wilcoxon signed rank tests. Unless otherwise noted, all tests were two-tailed and all plots with error bars were reported as mean ± SEM; all boxplots show the median (center line), interquartile range (box limits: 25th to 75th percentile), and whiskers (extending to the most extreme data points within 1.5× the interquartile range). Outliers beyond the whiskers are shown as individual points. Sample size was not predetermined using power analysis.

## Data availability

All behavioral and electrophysiological data will be made available upon publication.

